# Convergent and Divergent Cerebellar Alterations in 22q11.2 Copy Number Variants

**DOI:** 10.1101/2025.10.07.679846

**Authors:** Hoki Fung, Kathleen P. O’Hora, Rune Boen, Carolyn M. Amir, Charles H. Schleifer, Leila Kushan-Wells, Blake A. Miranda, Elizabeth Bondy, Paul J. Mathews, Carrie E. Bearden

## Abstract

**Background:** Individuals with copy number variants (CNVs) at 22q11.2 are at elevated risk for neurodevelopmental and psychiatric disorders, including autism spectrum disorder and intellectual disability. For psychosis, effects diverge, with 22q11.2 deletion (22qDel) conferring one of the highest known risks for schizophrenia, while duplication (22qDup) may be protective. Prior investigations of neurobiological mechanisms in 22q11.2 CNVs have predominantly focused on the cerebrum, whereas the cerebellum—a region increasingly recognized for its contributions to cognitive, affective, and social processes—remains underexplored and represents a promising target for investigation. Although alterations in cerebellar structure have been reported in 22qDel, they remain largely unexplored in 22qDup. This study provides the first characterization of regional cerebellar volumes in 22qDup and the first direct comparison of cerebellar structure across 22q11.2 CNVs, offering a unique opportunity to identify shared and distinct neurobiological mechanisms with implications for understanding cerebellar contributions to brain–behavior relationships in CNV carriers.

**Methods:** We analyzed 514 longitudinally collected structural Magnetic Resonance Imaging (MRI) scans of 111 individuals with 22qDel, 37 individuals with 22qDup, and 167 typically developing (TD) controls. Total and regional cerebellar volumes were quantified using ACAPULCO, a deep-learning-based parcellation method that segments the cerebellum into 28 subregions. Group differences in cerebellar volumes, as well as their associations with cognition, autism-related traits, and psychosis-risk symptoms, were examined using linear mixed-effects models. False discovery rate (FDR) correction was applied to control for multiple comparisons where appropriate.

**Results:** In relation to TD controls, cerebellar volumes were broadly reduced in 22qDel, whereas cerebellar alterations in 22qDup were more modest and variable. Regional analyses revealed both linear (22qDel < TD < 22qDup) and nonlinear (22qDel ≈ 22qDup < TD; 22qDel < 22qDup < TD) gene-dosage patterns, though not all reached significance. Vermis VII was significantly reduced in both CNVs but showed no relationship to behavioral differences. In contrast, reduction of Right Lobule VIIIA was associated with greater social impairment in 22qDup. Unlike TD controls, this region was not associated with IQ in 22qDup, suggesting CNV-specific alterations in cerebellar–behavior relationships.

**Conclusion:** These findings indicate that reciprocal CNVs at the 22q11.2 locus both affect cerebellar structure, yet their functional consequences diverge, suggesting overlapping but distinct pathways to clinical risk. They underscore the cerebellum’s multifaceted role in neurodevelopment and highlight the need for studies examining broader cognitive and socio-affective domains, as well as cerebellar–cortical connectivity, to clarify links to clinical outcomes.

## Introduction

Copy number variants (CNVs; i.e., deletions or duplications) at the 22q11.2 locus have a profound impact on neurodevelopment and clinical outcomes, encompassing a wide range of cognitive, psychiatric, and socio-affective symptoms (Fiksinski et al., 2023; McDonald-McGinn et al., 2015; Schneider et al., 2014). 22q11.2 deletion (22qDel), a ∼1.5-2.6 Megabase hemizygous deletion on the long arm of chromosome 22, is the most common contiguous gene deletion syndrome, occurring in approximately 1 in 3,700 live births (Olsen et al., 2018). Along with craniofacial abnormalities, congenital heart defects, and other medical conditions, 22qDel confers greatly increased likelihood of developmental neuropsychiatric disorders including autism spectrum disorder (ASD), intellectual disability (ID), attention deficit hyperactivity disorder (ADHD), and anxiety disorders (Schneider et al., 2014). Notably, 22qDel is one of the strongest known genetic risk factors for schizophrenia, with approximately 1 in 10 carriers of a 22qDel meeting criteria for schizophrenia (Provenzani et al., 2022) and approximately one-third exhibiting clinically significant sub-threshold psychosis symptoms (Jalbrzikowski et al., 2022; Weisman et al., 2017). The 22q11.2 duplication (22qDup) is the reciprocal of the 22qDel, occurring at the same locus and of comparable size (Ou et al., 2008). Although less well-characterized than 22qDel, the 22qDup may actually be more common, with an estimated prevalence of ∼1 in 1,600 live births (Olsen et al., 2018). The 22qDup confers similarly increased likelihood of ASD, ADHD, and anxiety disorders relative to the general population as 22qDel, but it does not confer the same elevated risk for schizophrenia (Olsen et al., 2018). In fact, 22qDup carriers may be less likely to develop schizophrenia (Marshall et al., 2017; Elliott Rees et al., 2014), making reciprocal 22q11.2 CNV carriers an intriguing model for studying biological underpinnings of psychosis risk. Deep phenotyping of structural brain features in 22q11.2 CNVs can thus further understanding of aberrant neurodevelopment and gene-brain-behavior relationships. Indeed, 22q11.2 CNVs have been associated with discernible alterations in cortical subcortical, and white matter structures (Ching et al., 2020; Kumar et al., 2023; Lin et al., 2017; Rogdaki et al., 2020; Schleifer et al., 2024; Seitz-Holland et al., 2021; Sun et al., 2020), often showing reciprocal effects between deletions and duplications. Specifically, 22qDel carriers exhibit smaller intracranial, gray, and white matter volume, on average, relative to controls (Ching et al., 2020; Lin et al., 2017; Rogdaki et al., 2020; Sun et al., 2020). In addition, 22qDel is associated with reduced cortical surface area but overall thicker cortices, along with smaller hippocampal volume, larger corpus callosum volume and higher fractional anisotropy assessed via diffusion weighted imaging, whereas 22qDup carriers show the opposite pattern (Lin et al., 2017; Schleifer et al., 2024; Seitz-Holland et al., 2021). These effects suggest that variation in copy number at the 22q11.2 locus contributes to the observed differences in brain structure.

The cerebellum contains up to 80% of all neurons in the human brain, despite only representing 10% of total brain mass (Azevedo et al., 2009). It is traditionally viewed as a motor structure, but receiving an increasing amount of focus as a critical hub for cognitive, affective, and social processing, with rapidly growing relevance across neuropsychiatric and neurodevelopmental disorders, including schizophrenia and ASD (Buckner, 2013). Neuroimaging studies have revealed a topographic organization of the cerebellum, with motor functions localized to the anterior lobe and part of lobules IV and VIII, cognitive functions to the posterior lobe—primarily lobule VII—and affective processes to the posterior vermis and adjacent regions (Buckner et al., 2011; King et al., 2019; Stoodley & Schmahmann, 2009). Reduced cerebellar grey matter volume has been consistently reported in idiopathic schizophrenia, with widespread reductions across multiple cerebellar lobules. These reductions are most pronounced in posterior regions linked to higher-order cognitive networks, while anterior, motor-related regions appear relatively less affected (Moberget et al., 2018). In contrast, findings in ASD are less consistent, with some studies reporting increased and others reporting reduced total cerebellar volume (Mapelli et al., 2022; Rodrigues et al., 2025). The inconsistencies likely reflect the heterogeneity and complexity of ASD.

Most cerebellar research on schizophrenia and ASD has focused on idiopathic cases. In contrast, the effects of 22q11.2 CNVs—genetically defined variants that modulate risk for both disorders—on cerebellar structure and function remain understudied, even though they provide a more uniform model for linking cerebellar structure to behavior than heterogeneous idiopathic samples. Lower overall cerebellar volume in 22qDel carriers has consistently been reported in cross-sectional studies, in childhood, adolescence, and adulthood (Bish et al., 2006; Campbell et al., 2006; Gothelf et al., 2007; Shashi et al., 2010; van Amelsvoort et al., 2004, 2001). More recent work by Schmitt et al. (2023) examined the cerebellum in 22qDel at a finer, lobular resolution and found significant volume reductions across all subregions (Schmitt et al., 2023). In contrast, no study to date has investigated the cerebellum in 22qDup carriers, so it is unknown if the same gene-dosage effect observed in other brain regions is present in the cerebellum. Addressing this gap also provides a unique opportunity to examine whether common structural alterations in 22qDel and 22qDup relate to distinct patterns of symptomatology across the two CNVs.

Here, we investigate volumetric differences in the cerebellum and its subregions across 22q11.2 CNV carriers and typically developing (TD) controls. Drawing on prior literature, we hypothesize that: 1) 22q11.2 CNVs will exhibit positive gene-dosage effects on total cerebellar volume, with 22qDel carriers showing decreased volume and 22qDup carriers showing increased volume relative to controls (22qDel < TD < 22qDup), and 2) similar positive gene-dosage effects across cerebellar subregions. To clarify the functional relevance of these volumetric alterations, we further test whether subregions showing gene-dosage dependent volume differences are related to clinical outcomes relevant to 22q11.2 CNVs, including cognition, psychosis risk, and autistic traits, and whether these associations differ from those observed in TD controls. We address these aims through collection and analysis of the largest sample of reciprocal 22q11.2 CNVs to date and application of ACAPULCO, a novel cerebellar parcellation method (Han et al., 2020), to structural Magnetic Resonance Imaging (MRI) data.

## Methods

### Participants and Study Design

In this study, we collected 524 longitudinal structural MRI scans from 317 unique individuals. The final analysis included 514 scans from 315 participants: 111 with 22qDel, 37 with 22qDup, and 167 TD controls. Participants with molecularly confirmed 22qDel or 22qDup were recruited as part of an ongoing longitudinal study at the University of California, Los Angeles (UCLA). Recruitment occurred through clinical referrals, community outreach, and research registries. TD controls were recruited from the same communities as CNV carriers. All participants underwent cognitive testing, clinical assessments, and structural MRI scanning across one to six annual visits (mean number of visits per participant = 1.63). TD controls were screened to exclude any personal or family history of psychosis, major neurological disorders, or known genetic conditions. Additional exclusion criteria for all participants included history of significant head trauma, contraindications to MRI, or severe sensorimotor impairments precluding valid assessment. Written informed consent (or assent with parental consent for minors) was obtained from all participants. All procedures were approved by the Institutional Review Board at UCLA. Further details on inclusion and exclusion criteria, recruitment procedures, scanning protocol, and clinical measures have been reported previously (Jalbrzikowski et al., 2022; Schleifer et al., 2024).

### Behavioral and Clinical Measures

#### Cognitive Ability

Overall cognitive ability was assessed using age-appropriate standardized measures of IQ, the *Wechsler Abbreviated Scale of Intelligence* (WASI; (Wechsler, 1999)) or the *Wechsler Abbreviated Scale of Intelligence – Second Edition* (WASI-II; (Wechsler, 2011)). Full Scale IQ (FSIQ) scores were calculated based on established scoring procedures (Wechsler, 1999). Testing was conducted by trained research staff under the supervision of licensed psychologists.

#### Autism-related Traits

Autism-related traits were measured using the *Social Responsiveness Scale 2nd edition* (SRS; (Constantino & Gruber, 2005)), a well-validated parent-report questionnaire that quantifies impairments in reciprocal social behavior. SRS total scores were used as the primary outcome variable to capture dimensional variation in autism-related symptomatology across the sample.

#### Psychosis Risk Symptoms

Psychosis risk symptoms were assessed in participants over the age of ten using the *Structured Interview for Psychosis-Risk Syndromes* (SIPS; (McGlashan et al., 2001)), administered by trained Master’s-level clinicians with supervision from a licensed clinical psychologist. The total positive symptom score was used as the primary outcome measure. It sums severity ratings of five items assessing positive symptoms: unusual thought content/delusional ideas, suspiciousness/persecutory ideas, grandiose ideas, perceptual abnormalities/hallucinations, and disorganized communication.

### MRI Acquisition

Structural MRI data were collected at UCLA using three 3T Siemens MRI scanners: a Trio system at the Ahmanson-Lovelace Brain Mapping Center (BMC), and the Trio and Prisma systems at the Staglin Center for Cognitive Neuroscience (CCN). High-resolution T1-weighted images were acquired using standardized magnetization-prepared rapid gradient echo (MPRAGE) sequences (Mugler & Brookeman, 1990) using comparable parameters across scanners (TR = 2,300–2,400 ms, TE = 2.22–2.89 ms, TI = 900–1,060 ms, flip angle = 8–9°, voxel size = 1.0 × 1.0 × 1.2 mm or 0.8 mm^3^ isotropic, and field of view = 240 × 256 mm^2^). Data were subsequently harmonized to reduce scanner-related variability, as described in the Data Harmonization section.

### T1 Image Processing

Prior to processing, all T1-weighted structural images were visually inspected for motion, artifacts, and acquisition errors. High-quality raw images were then used as input to ACAPULCO (version 3.1; (Han et al., 2020)) for cerebellar segmentation. ACAPULCO is a deep learning-based parcellation tool that divides the cerebellum into 28 anatomically defined subregions and includes its own internal preprocessing pipeline. The ACAPULCO pipeline was run in a containerized environment to ensure consistency across systems. Following segmentation, outputs were visually reviewed for quality, and scans with failed or poor parcellations were excluded from further analysis. Estimated total intracranial volume (eTIV) was derived from the T1-weighted images using FreeSurfer (versions 5.3.0 or 6.0.0; (Fischl, 2012)), and used as a covariate in all volumetric analyses.

### Data Harmonization

To address scanner-related variability in volume measures, we applied Longitudinal ComBat (LongComBat) harmonization (Beer et al., 2020) to eTIV and cerebellar volumes. LongComBat adjusts for scanner-related effects in multi-site longitudinal data while aiming to preserve biologically meaningful variance (Beer et al., 2020).

### Statistical Analysis

#### Cerebellar Volume

To examine group differences in cerebellar volume, we fit linear mixed-effects models. Separate models were run for total cerebellar volume and for each of 28 anatomically defined cerebellar subregions (see Supplementary Table S2 for the full list). Each model included fixed effects for group (TD, 22qDel, 22qDup), sex, age (mean centered linear and quadratic terms), and eTIV, with a random intercept for subject to account for repeated measures across visits. TD controls were used as the reference group for all models unless otherwise specified. To account for multiple comparisons, False Discovery Rate (FDR) correction was applied across the 28 regional volume models.

#### Brain–Behavior Associations

To examine the relationship between cerebellar structure and neurobehavioral outcomes, we tested associations between regional cerebellar volumes and **1)** IQ, **2)** SRS total scores, and **3)** SIPS total positive symptom scores using all available data across visits, including repeated measures where present. To limit the number of statistical comparisons in this exploratory analysis and focus on regions most likely to show clinically meaningful associations, brain-behavior associations were tested only in cerebellar regions that showed significant volumetric differences in both 22qDel and 22qDup groups relative to controls (*p*s < 0.05).

### IQ and SRS

To evaluate whether the relationship between regional cerebellar volumes and behavioral outcomes differed across groups, we fit separate linear mixed-effects models for each *a priori*-defined region-outcome pair. The behavioral outcome (IQ or SRS) was the dependent variable, and the primary predictor of interest was the *interaction* between volume and group (TD, 22qDel, 22qDup). All models included fixed effects for sex, age (mean centered linear and quadratic terms), and eTIV, with a random intercept for subject to account for repeated measures. All analyses were hypothesis-driven and restricted to regions that showed convergent reductions in both 22qDel and 22qDup groups relative to TD controls, given prior evidence of convergent alterations in cognition and social functioning in 22q11.2 CNVs (i.e., lower IQ and elevated ASD traits; (Lin et al., 2020)). As analyses were confined to these theoretically motivated regions, rather than conducted across the full set of cerebellar regions, FDR correction was not applied.

### SIPS Positive Symptoms

To align with the observed psychosis risk behavioral pattern (i.e., elevated risk in 22qDel carriers and lower risk in 22qDup carriers (Marshall et al., 2017; Elliott Rees et al., 2014)), we examined whether the relationship between regional cerebellar volume and SIPS total positive symptom scores differed across groups in regions that showed corresponding volumetric differences (22qDel < TD < 22qDup). The modeling framework and covariates were the same as those used for the IQ and SRS analyses, with the exception that SIPS total positive symptom scores were log-transformed to address substantial zero inflation and a right-skewed distribution (O’Hora et al., 2024). As floor effects were observed in the TD control group, we additionally examined within-group associations for 22qDel and 22qDup participants, using the same covariates as the across-group analysis.

#### Data Scaling and Visualization

All statistical models reported in text and tables were fitted using raw, unstandardized values to preserve interpretability in native units (e.g., mm³ for brain volumes, points for behavioral scores). For visualization purposes, some figures (e.g., forest plots, flatmaps) display standardized values to facilitate comparison across regions and groups. Please refer to individual figure legends for details.

## Results

### Participant Characteristics

The full sample included 317 unique participants and 524 MRI sessions. Following parcellation quality control, 10 scans were excluded due to poor-quality cerebellar segmentations, resulting in a final analytic sample of 514 sessions from 315 participants.

Baseline demographic and behavioral characteristics of the study sample, along with group comparison statistics, are summarized in Table 1. Briefly, the three groups (TD controls, 22qDel, and 22qDup) were comparable in sex and age distributions at baseline. IQ was lower in both CNV groups (22qDel < 22qDup < TD). SRS total scores, reflecting autism-related traits, were elevated in both CNV groups relative to controls (TD < 22qDel ≈ 22qDup). SIPS total positive symptom scores, reflecting psychosis severity, were higher in 22qDel relative to both TD controls and 22qDup, with no significant difference between TD controls and 22qDup (TD ≈ 22qDup < 22qDel). eTIV was lower in 22qDel participants compared to both TD control and 22qDup groups (22qDel < TD ≈ 22qDup). These patterns are consistent with a previous cross-sectional characterization of phenotypic features of 22q11.2 CNVs in a subset of this cohort (Lin et al., 2020). See Table 1 for full statistical results.

**Table 1.**
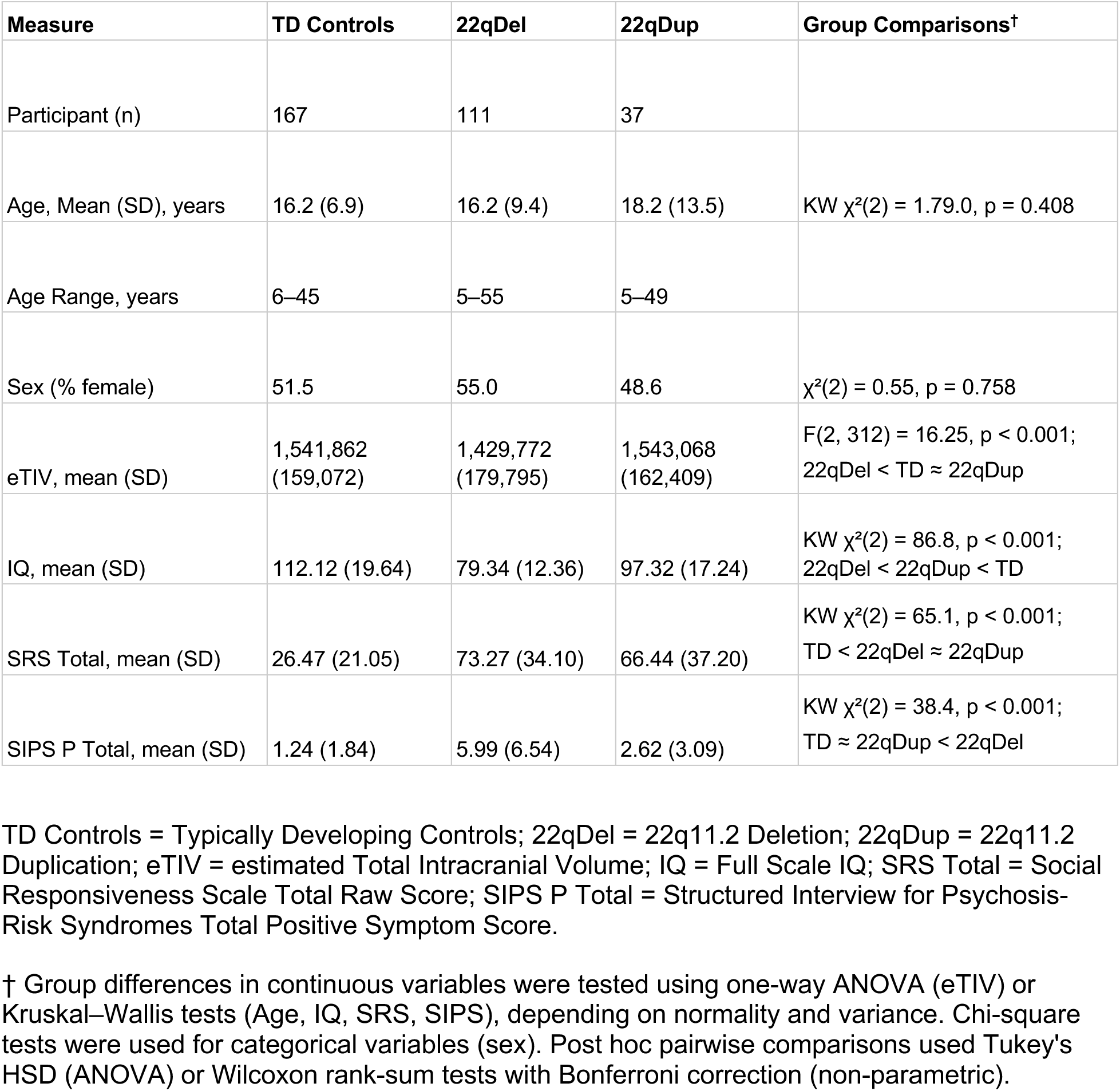
Participant Characteristics at Baseline by Group.

| Measure | TD Controls | 22qDel | 22qDup | Group Comparisons<sup>†</sup> |
| --- | --- | --- | --- | --- |
| Participant (n) | 167 | 111 | 37 |  |
| Age, Mean (SD), years | 16.2 (6.9) | 16.2 (9.4) | 18.2 (13.5) | KW $\chi^2(2) = 1.79.0$, $p = 0.408$ |
| Age Range, years | 6–45 | 5–55 | 5–49 |  |
| Sex (% female) | 51.5 | 55.0 | 48.6 | $\chi^2(2) = 0.55$, $p = 0.758$ |
| eTIV, mean (SD) | 1,541,862<br>(159,072) | 1,429,772<br>(179,795) | 1,543,068<br>(162,409) | $F(2, 312) = 16.25$, $p < 0.001$;<br>22qDel < TD ≈ 22qDup |
| IQ, mean (SD) | 112.12 (19.64) | 79.34 (12.36) | 97.32 (17.24) | KW $\chi^2(2) = 86.8$, $p < 0.001$;<br>22qDel < 22qDup < TD |
| SRS Total, mean (SD) | 26.47 (21.05) | 73.27 (34.10) | 66.44 (37.20) | KW $\chi^2(2) = 65.1$, $p < 0.001$;<br>TD < 22qDel ≈ 22qDup |
| SIPS P Total, mean (SD) | 1.24 (1.84) | 5.99 (6.54) | 2.62 (3.09) | KW $\chi^2(2) = 38.4$, $p < 0.001$;<br>TD ≈ 22qDup < 22qDel |

TD Controls = Typically Developing Controls; 22qDel = 22q11.2 Deletion; 22qDup = 22q11.2 Duplication; eTIV = estimated Total Intracranial Volume; IQ = Full Scale IQ; SRS Total = Social Responsiveness Scale Total Raw Score; SIPS P Total = Structured Interview for Psychosis-Risk Syndromes Total Positive Symptom Score.
<sup>†</sup> Group differences in continuous variables were tested using one-way ANOVA (eTIV) or Kruskal–Wallis tests (Age, IQ, SRS, SIPS), depending on normality and variance. Chi-square tests were used for categorical variables (sex). Post hoc pairwise comparisons used Tukey's HSD (ANOVA) or Wilcoxon rank-sum tests with Bonferroni correction (non-parametric).

### Total Cerebellar Volume

A linear mixed-effects model revealed that individuals with 22qDel had significantly smaller total cerebellar volume compared to TD controls (*B* = −16,072.03, *p* < 0.001), while in contrast 22qDup displayed no significant difference to TD controls (*B* = 1,579.76, *p* = 0.46; see Supplementary Table S1 for full model output). However, the group means followed a consistent directional trend, with 22qDel showing the lowest average volume, TD controls intermediate, and 22qDup the highest.

### Regional Cerebellar Volume

To determine whether overall group patterns in total cerebellar volume reflected localized effects, we conducted region-wise analyses across 28 anatomically defined regions. Figure 1 summarizes standardized beta coefficients and 95% confidence intervals for both 22qDel and 22qDup groups, across all cerebellar regions.

**Figure 1.**
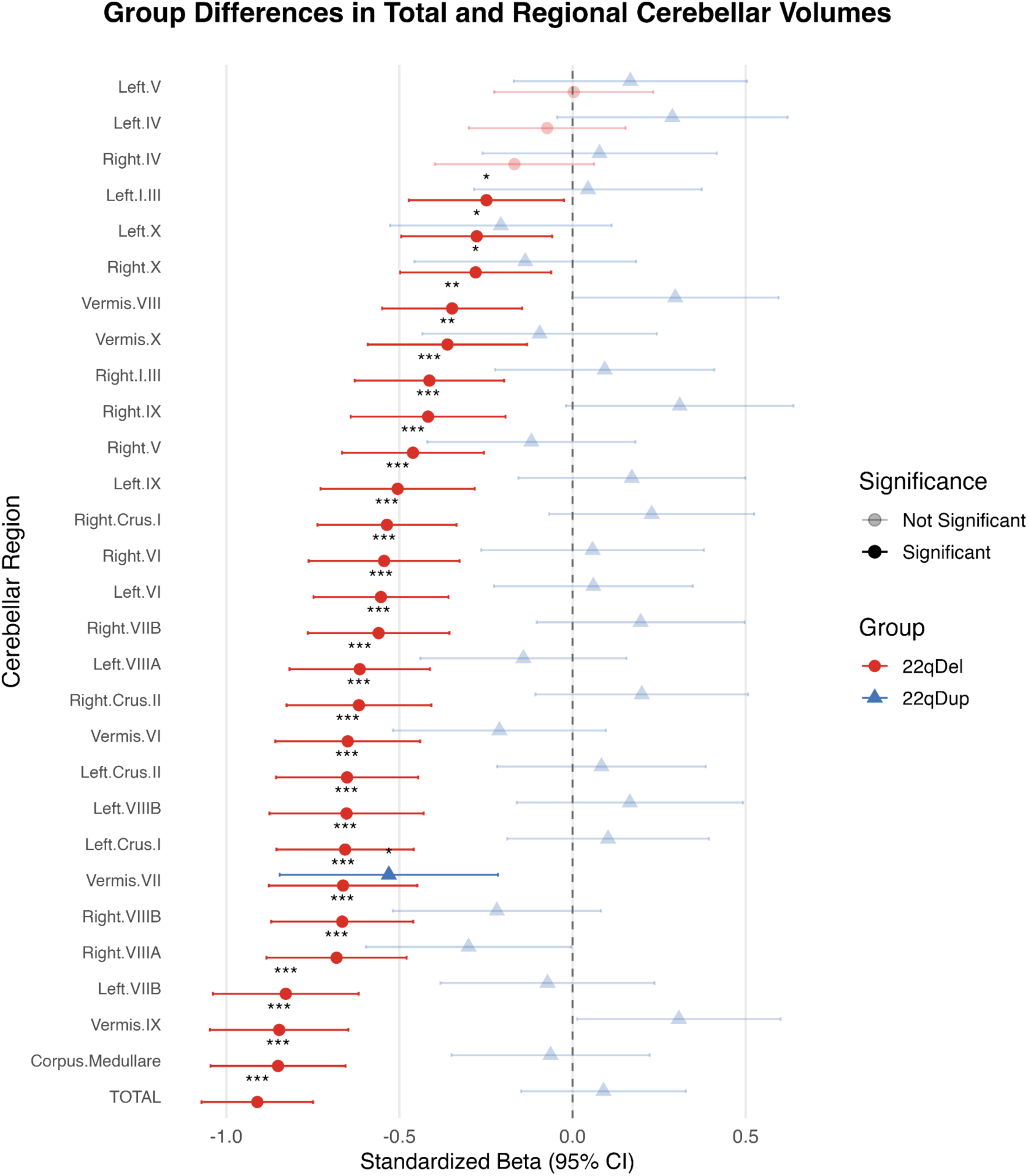
Group differences in total and regional cerebellar volumes. Standardized beta coefficients and 95% confidence intervals are shown for 22qDel and 22qDup groups relative to typically developing controls. Results were derived from linear mixed-effects models controlling for sex, mean-centered age (linear and quadratic), and estimated intracranial volume (eTIV), with a subject-level random intercept to account for repeated measures. Asterisks indicate significance after FDR correction for regional volumes: *q* < .05 (*), *q* < .01 (**), *q* < .001 (***). TOTAL refers to total cerebellar volume and is shown with uncorrected p-value, as it was not included in the FDR correction.

Individuals with 22qDel showed widespread cerebellar volume reductions relative to TD controls (Figure 2A). Specifically, 25 out of 28 regions (except bilateral Lobule IV and Left Lobule V) were significantly smaller in the 22qDel group after FDR correction (*q* < 0.05; Figure 2A), indicating diffuse cerebellar involvement in 22q11.2 deletion carriers. Among these, the largest reductions were observed in the **Corpus Medullare** (*B* = −2268.34, *p* < 0.001), **Vermis IX** (*B* = −168.46, *p* < 0.001), and **Left Lobule VIIB** (*B* = −1178.50, *p* < 0.001). See Supplementary Table S2 for full model results.

**Figure 2.**
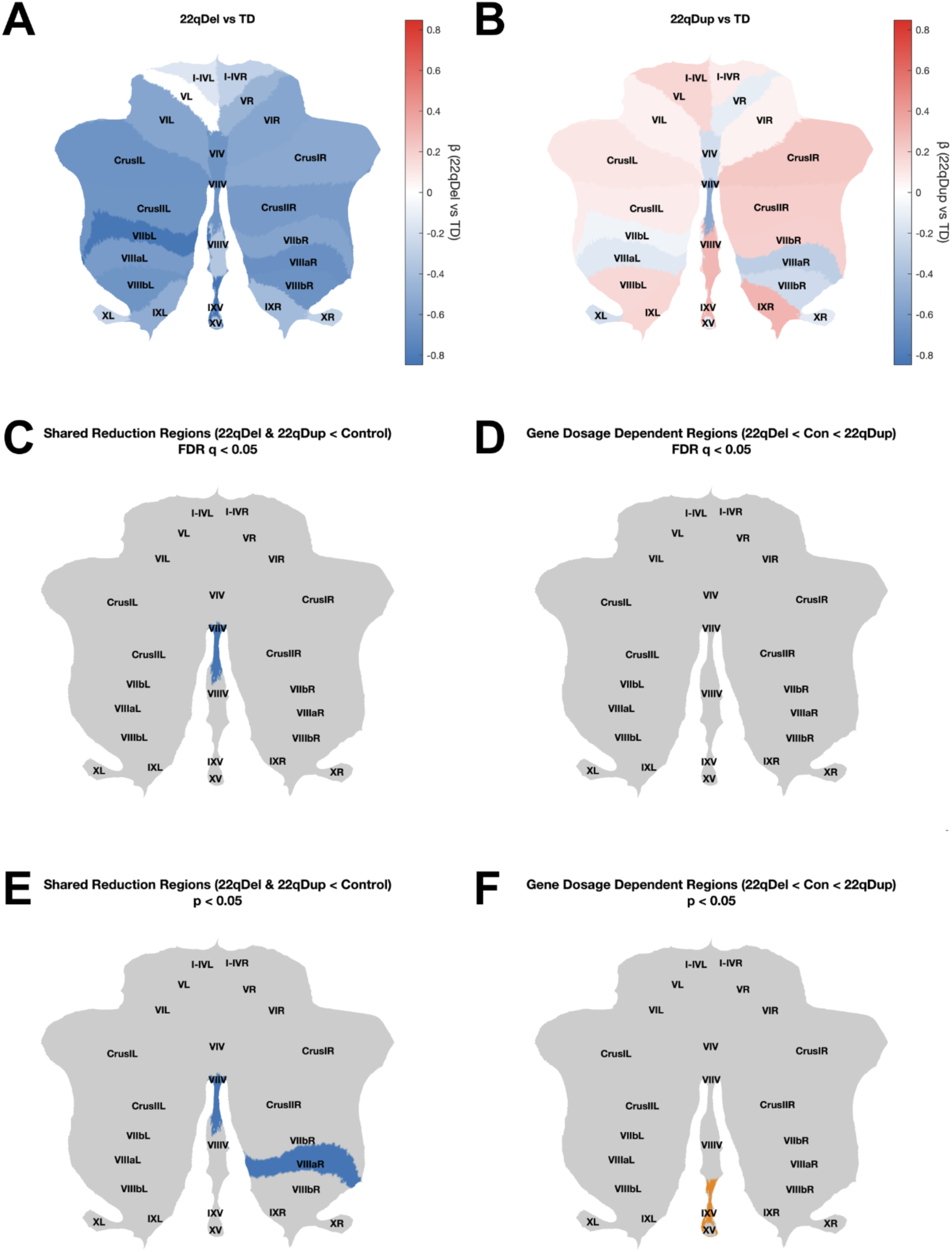
Flatmap visualizations of cerebellar volume differences across 22q11.2 CNVs. **Panel A:** Flatmap indicating widespread cerebellar volume reductions in 22qDel compared to TD, particularly in midline and lateral posterior regions. **Panel B:** Flatmap indicating more subtle and heterogeneous volume differences in 22qDup compared to TD, including both increases and reductions across subregions. **Panel C:** Binary map highlighting regions with convergent volume reductions in 22qDel and 22qDup at FDR q < 0.05 (Vermis VII). **Panel D:** Binary map highlighting regions with monotonic gene dosage effects (22qDel < TD < 22qDup) at FDR q < 0.05 (no regions). **Panel E:** Binary map highlighting regions with convergent volume reductions in 22qDel and 22qDup at uncorrected p < 0.05 (Vermis VII, Right Lobule VIIIA). **Panel F:** Binary map highlighting regions with monotonic gene dosage effects (22qDel < TD < 22qDup) at uncorrected p < 0.05 (Vermis IX). Regional volumes were derived using the ACAPULCO cerebellar parcellation and mapped to SUIT-defined anatomical regions for visualization. Beta values reflect standardized group effects from linear mixed-effects models, displayed without p-value thresholding on a symmetric scale (blue = negative, red = positive). To align with SUIT’s anatomical scheme, beta values of Lobule I–III and Lobule IV from ACAPULCO were averaged to create a single Lobule I–IV region. The corpus medullare was not visualized in these maps. Binary maps show regions where both 22qDel and 22qDup differed significantly from TD at the indicated threshold; gray regions indicate regions that did not meet this criterion. See Supplementary Table S2 for full model results.

Unlike the consistently smaller cerebellar volumes observed in 22qDel carriers, individuals with 22qDup showed a more heterogeneous pattern of regional volume differences, with beta estimates indicating larger volumes in 17 regions and smaller volumes in 11 regions, relative to TD controls (Figure 2B; Supplementary Table S2 for full model results). However, these effects were generally smaller, with several regions reaching nominal significance at *p* < 0.05, but only one region surviving FDR correction (*q* < 0.05). Specifically, **Vermis VII** (*B* = −100.05, *p* < 0.001, *q* < 0.05) was smaller in 22qDup compared with TD controls. At the nominal threshold (*p* < 0.05), **Right Lobule VIIIA** (*B* = −358.09, *p* = 0.047, *q* = 0.36) was smaller and **Vermis IX** (*B* = 61.02, *p* = 0.041, *q* = 0.36) was larger in 22qDup relative to TD controls.

Direct comparisons between the two CNV groups, through reparameterizing the original models with 22qDel as the reference group, revealed that 22qDup carriers had significantly larger volumes than 22qDel carriers in the vast majority of regions (20 of 28; Figure S1; Table S3).

Next, from the TD control-referenced comparisons, we identified regions that showed convergently smaller volumes in both CNV groups vs. those consistent with a gene-dosage dependent effect (22qDel < TD < 22qDup). Convergent or gene dosage effects were defined as regions where both 22qDel and 22qDup differed significantly from TD controls at the same threshold. At the FDR-corrected level (*q* < 0.05), **Vermis VII** was smaller in both groups (Figure 2C). At the nominal level (*p* < 0.05), **Right Lobule VIIIA** was also smaller in both groups (Figure 2E), and **Vermis IX** followed the gene dosage pattern (Figure 2F).

To further characterize the convergently smaller regions, we revisited the direct 22qDel vs 22qDup comparison (Figure S1; Table S3) to evaluate whether both CNV groups exhibited comparably smaller volumes relative to TD controls (22qDel ≈ 22qDup < TD) or whether they followed a gradient (22qDel < 22qDup < TD). Vermis VII volumes did not differ significantly between 22qDel and 22qDup (*B* = 24.93, *p* = 0.43). In Right Lobule VIIIA, however, the volume was significantly higher in 22qDup than in 22qDel (*B* = 454.53, *p* = 0.02), indicating a gradient. Together with Vermis IX, these three regions were selected for exploratory brain–behavior analyses based on the *a priori* strategy described in the Methods, which targeted cerebellar regions showing alterations in both CNV groups relative to TD controls.

### Associations Between Cerebellar Volume and Neurobehavioral Outcomes

#### Cognitive Ability (IQ)

In the two cerebellar regions—Vermis VII and Right Lobule VIIIA—that were significantly smaller in both 22qDel and 22qDup compared to controls (Figure 3A–B), we examined the relationship between regional cerebellar volume and Full Scale IQ across groups using linear mixed-effects models with a Group x Volume interaction term. In TD controls, volumes in both Vermis VII (*B* = 0.017, *p* = 0.038) and Right Lobule VIIIA (*B* = 0.0036, *p* = 0.003) were significantly positively associated with IQ, reflecting robust cerebellar volume-IQ associations. In 22qDel, however, the positive relationships with IQ were weaker and did not reach significance for either Vermis VII (*B* = 0.0036, *p* = 0.63) or Right Lobule VIIIA (*B* = 0.0015, *p* = 0.095). However, Group x Volume interaction terms were non-significant, indicating no significant difference in volume-IQ associations between 22qDel and TD controls. In 22qDup, the volume-IQ association in Vermis VII was also not significant (*B* = 0.0052, *p* = 0.63) and did not differ significantly from TD controls (interaction *B* = −0.012, *p* = 0.38). In Right Lobule VIIIA, however, the volume-IQ association differed significantly from TD controls (interaction *B* = −0.0044, *p* = 0.038; Figure 3B).

**Figure 3.**
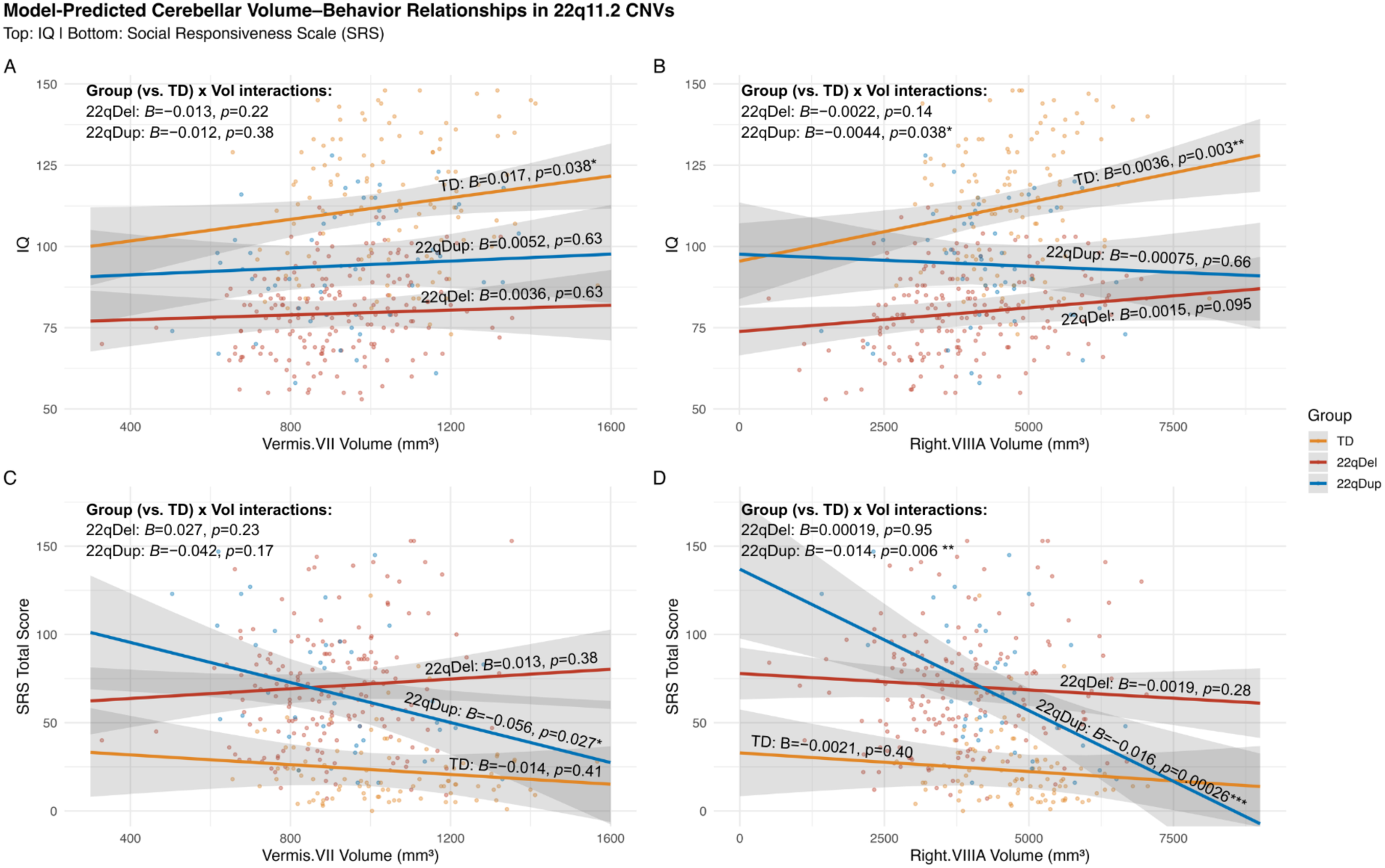
Model-predicted relationships between cerebellar volume and behavior in regions convergently reduced in 22q11.2 CNVs. Each panel shows the modeled association between regional cerebellar volume and behavioral outcome by group (TD, 22qDel, 22qDup). Solid lines show group-specific fixed-effects predictions from linear mixed-effects models that included cerebellar volume, group, and their interaction, with age (mean-centered linear and quadratic), sex, and eTIV as covariates, and a random intercept for subject. Shaded ribbons denote 95% confidence intervals around the predicted means. Semi-transparent points show observed behavioral outcome values plotted against LongComBat-harmonized cerebellar volumes. Displayed alongside each line are within-group slope estimates with corresponding p-values, and interaction p-values are shown in the upper left corner of each panel. Panel A: Volume of Vermis VII vs. IQ. Panel B: Volume of Right Lobule VIIIA vs. IQ. Panel C: Volume of Vermis VII vs. SRS total score. Panel D: Volume of Right Lobule VIIIA vs. SRS total score.

#### Autism Traits (SRS)

Given the similar elevation of ASD traits in both 22q11.2 CNV groups, this analysis focused on Vermis VII and Right Lobule VIIIA, the regions that showed volume reductions in both 22qDel and 22qDup relative to controls (Figure 3C–D). In controls, neither region showed a significant association with SRS total scores, suggesting an absence of cerebellar structure–social function coupling in typical development (Vermis VII: *B* = −0.014, *p* = 0.41; Right Lobule VIIIA: *B* = −0.0021, *p* = 0.40). However, within the 22qDup group, smaller volumes in both Vermis VII (*B* = −0.056, *p* = 0.027) and Right Lobule VIIIA (*B* = −0.016, p < 0.001) were significantly associated with greater social impairment. While the volume–SRS association in Vermis VII did not significantly differ from TD controls, a robust interaction was observed in Right Lobule VIIIA (Interaction *B* = −0.014, *p* = 0.006; Figure 3D). This pattern was not observed in 22qDel, however (Vermis VII: interaction *B* = 0.027, *p* = 0.23; Right Lobule VIIIA: interaction *B* = 0.00019, *p* = 0.95).

Together, these findings point to a group-specific alteration in cerebellar–social behavior coupling in 22qDup, where smaller volume in Right Lobule VIIIA is associated with elevated autism-related traits, a pattern not observed in 22qDel or TD controls.

#### Psychosis-Risk Symptoms (SIPS)

Guided by our *a priori* rationale described in the Methods, we tested whether volume in Vermis IX, a region showing gene-dosage-dependent differences, was associated with SIPS positive symptom severity, and whether this association varied across groups. Vermis IX volume was not significantly related to SIPS total positive symptom scores in any group, and no Group x Volume interaction was observed. Within-group analyses restricted to 22qDel or 22qDup yielded similar null results.

## Discussion

This study provides the first direct comparison of regional cerebellar volumes across 22q11.2 CNVs. Using high-resolution T1-weighted MRI and a cutting-edge deep-learning cerebellar parcellation pipeline, we uncovered distinct and convergent patterns of cerebellar morphology across 22q11.2 CNV carriers. 22q11.2 duplication carriers had cerebellar volumes largely comparable to controls, yet exhibited a significant reduction in Vermis VII and nominal trends toward slightly enlarged Vermis IX and smaller Right Lobule VIIIA. In contrast, consistent with previous reports, 22q11.2 deletion carriers displayed robust and widespread reductions in both total and regional cerebellar volume. Notably, Vermis VII was similarly reduced in both CNV groups relative to controls, highlighting a shared locus of structural vulnerability. Although this convergent reduction was not linked to IQ or autism-related traits, Right Lobule VIIIA was notable for its functional relevance: in 22qDup carriers only, smaller volumes were associated with greater social impairment. This points to a potential duplication-specific alteration in cerebellar-social behavior coupling. To our knowledge, this is the first study to characterize regional cerebellar volumetric alterations in individuals with 22q11.2 duplications, a recently characterized CNV associated with high risk for autism and possibly protective against schizophrenia (Marshall et al., 2017; E. Rees et al., 2014; Shanta et al., 2025).

### Widespread Cerebellar Volume Reduction in 22q11.2 Deletion

Our results are largely consistent with previously reported findings of widespread cerebellar volume reductions in 22qDel ((Schmitt et al., 2023); N=79). In our larger, independent sample of 22qDel carriers, nearly all cerebellar subregions—with the exception of bilateral Lobule IV and Left Lobule V, both located in the anterior lobe of the cerebellum—were significantly smaller in 22qDel carriers relative to TD controls, highlighting diffuse structural vulnerability across both vermal and hemispheric lobules. These results largely replicate and extend prior evidence of global cerebellar hypoplasia in 22qDel (van Amelsvoort et al., 2004, 2001) and broader psychosis-related populations, including individuals with idiopathic schizophrenia and youth at ultra-high risk of psychosis (Bottmer et al., 2005; Dean et al., 2014; Moberget et al., 2018). These converging lines of evidence highlight the cerebellum as a key locus of neurodevelopmental disruption in 22qDel and behaviorally defined psychosis-risk populations.

Notably, the reduction in overall cerebellar volume in 22qDel is more pronounced (*d* = −0.91) than the previously reported reductions in neocortical gray matter (estimated at *d* ≈ −0.55) and white matter (estimated at d ≈ −0.88), both derived from values reported in (Lin et al., 2017). This pattern may reflect differential disruption of cerebral and cerebellar development across different stages of neurodevelopment, as these structures follow distinct spatiotemporal developmental trajectories. For instance, whereas neocortical expansion is primarily driven by radial glia cells in the ventricular and subventricular zone during early and mid-gestation (Nowakowski et al., 2016), cerebellar expansion is driven by radial glial, granule and Purkinje cell progenitors in the ventricular zone, rhombic lip and external granule layer during late gestation and postnatal periods (Leto et al., 2016; Volpe, 2009). Moreover, during late gestation, the cerebellum develops more rapidly than the cerebrum, with MRI studies showing a ∼3.5– 4-fold increase in cerebellar volume compared to only a 2–2.5-fold increase in cerebral volume during the same period (Limperopoulos et al., 2005). As such, reduced cerebellar volume may reflect particular disruption of progenitor cell production during late gestation in 22qDel (Farini et al., 2021), together with cortical reductions that likely arise from earlier neurodevelopmental processes, highlighting how distinct developmental windows contribute to differential vulnerability in 22qDel.

### Modest Cerebellar Alterations in 22q11.2 Duplication

In contrast to the widespread and robust reductions observed in 22qDel carriers, cerebellar alterations in 22qDup were comparatively modest and variable in direction. A significant reduction was observed in Vermis VII, alongside nominal trends toward enlargement of Vermis IX and reduction of Right Lobule VIIIA. The absence of a group difference in total cerebellar volume likely reflects this variable pattern, with local increases and decreases balancing out at the global level. This pattern suggests that duplication does not simply mirror the effects of deletion, but rather exerts smaller and possibly region-specific influences on cerebellar development. This underscores the importance of anatomically-detailed approaches that move beyond global measures to capture region-specific effects of genetic variation on brain structure.

As the first study to comprehensively characterize cerebellar morphology in 22qDup carriers using high-resolution, lobule-specific parcellation, there is limited prior literature to contextualize our results; however, these findings broadly align with emerging evidence from cortical and subcortical studies demonstrating that while certain brain measures (e.g. cortical thickness) scale proportionally with 22q11.2 gene dosage, others (e.g. total surface area, hippocampal volume) display asymmetry where the magnitude of reduction in 22qDel was greater than the increases observed in 22qDup (Lin et al., 2017; Schleifer et al., 2024). This pattern is also consistent with broader trends in the CNV literature, where duplications typically exert subtler and more variable effects than deletions, reflecting their generally lower penetrance across neuroanatomical and clinical phenotypes (Männik et al., 2015).

### Regional Cerebellar Alterations Across 22q11.2 CNVs

Vermis VII emerged as a robust locus of shared vulnerability, with both 22qDel and 22qDup carriers exhibiting significantly reduced volume relative to typically developing controls despite the opposing directions of 22q11.2 copy number variation. Right Lobule VIIIA showed a graded reduction (22qDel < 22qDup < TD) and Vermis IX a trend toward a positive dosage pattern, though these did not survive correction and may reflect subtler duplication effects. These findings highlight that cerebellar development is influenced by both linear gene-dosage relationships, where deletions and duplications have reciprocal effects, and non-linear patterns in which both deviate in the same direction—similar to prior reports of total gray and white matter volumes (Lin et al., 2017), although in that study the 22qDup versus TD controls difference was not significant. Vermis VII, part of the posterior “limbic cerebellum,” supports emotional regulation and higher-order cognitive functions (Jacobi et al., 2021; Schmahmann et al., 2021; Stoodley & Schmahmann, 2009, 2010), and reductions here align with reports of posterior vermis hypoplasia in autism and other neurodevelopmental conditions (Crucitti et al., 2020; Mapelli et al., 2022; Rodrigues et al., 2025). Right Lobule VIIIA contributes to sensorimotor, visual working-memory, and attentional processes (Brissenden et al., 2016, 2021, 2018; Cakar et al., 2024). It has also been implicated in ASD, with reduced gray matter volume correlating with greater social interaction symptom severity (D’Mello & Stoodley, 2015). Vermis IX is involved in vestibular (Laurens, 2022) and cardiovascular regulation (Bradley et al., 1991; La Noce et al., 1991) and altered connectivity in mood disorders (Saleem et al., 2023), underscoring the broad functional significance of these cerebellar regions in the context of 22q11.2 gene dosage.

### Dissociation Between Anatomical Vulnerability and Brain-Behavioral Relevance

When examining cerebellar volume–behavior relationships, the most robust deviations from TD controls emerged in 22qDup carriers. In this group, smaller Right Lobule VIIIA volume was associated with greater social impairment, a relationship not seen in controls, while the positive volume–IQ association observed in controls was absent. In contrast, no significant associations with cognition or ASD traits were detected in 22qDel carriers, despite pronounced reductions in the same region. These findings point to 22qDup-specific alterations in cerebellar structure–function coupling, though findings should be viewed as preliminary until replicated, given the modest sample size. Right Lobule VIIIA, a region involved in sensorimotor, working memory, and attentional processes (Brissenden et al., 2016, 2021, 2018; Cakar et al., 2024), may therefore play a distinct role in shaping cerebellar structure–behavior relationships in 22qDup.

By contrast, Vermis VII—despite being the most consistent locus of convergent anatomical reduction in both 22qDel and 22qDup—did not show associations with cognitive ability or autism-related traits that differed significantly from TD controls. This is noteworthy given that posterior vermal regions are involved in affective and social functioning and have often been implicated in autism and related conditions (Crucitti et al., 2020; Mapelli et al., 2022; Rodrigues et al., 2025). Future work in larger samples, incorporating additional cognitive and autism-related dimensions (including sensorimotor and attentional processes) and examining cerebellar functional connectivity with the broader brain, will be critical to clarifying these structure–function relationships.

### Limitations

Several limitations should be noted. Although our sample is the largest to examine cerebellar morphology in 22q11.2 CNVs, statistical power remains limited for brain-behavior analyses, especially for duplication carriers. Endorsement of positive psychosis risk symptoms was low in 22qDup and TD controls, as expected, and our relatively young 22qDel sample included fewer participants with overt psychosis compared with other clinically ascertained samples. These constraints may have impacted our ability to detect a linear gene-dosage effect in total or regional cerebellar volumes, as well as associations between cerebellar structure and psychosis-risk symptoms.

### Implications and Future Directions

Using a genetics-first approach, we find that both 22qDel and 22qDup carriers—each at elevated ASD risk—share a focal cerebellar vulnerability, with Vermis VII emerging as the most consistently reduced region. This converges with idiopathic ASD findings, positioning the cerebellum as a common substrate across genetic and non-genetic risk. Nominal effects in Right Lobule VIIIA and Vermis IX reveal regional heterogeneity, and smaller Right VIIIA volume tracks with greater social impairment in 22qDup, suggesting that behavioral impact can arise outside classic affective hubs. High-resolution lobular parcellation was key to detecting these region-specific effects, emphasizing the need for fine-grained brain–behavior mapping across CNV types.

Future studies using functional parcellation approaches may provide complementary insights into cerebellar volume differences in 22q11.2 CNVs beyond anatomical boundaries. Larger samples combining cerebellar functional connectivity, transcriptomics, and detailed behavioral phenotyping will also be critical. Finally, direct comparisons with other high-risk groups and neuropsychiatric CNVs (e.g., 3q29 and 16p11.2 CNVs) will further clarify shared and unique cerebellar mechanisms across neurodevelopmental disorders.

## Author Contributions

**H.F.:** Conceptualization, Methodology, Data Curation, Formal Analysis, Software, Validation, Visualization, Writing – Original Draft, Writing – Review & Editing. **K.P.O.:** Methodology, Data curation, Writing – Original Draft, Writing – Review & Editing. **R.B.:** Methodology, Writing – Original Draft, Writing – Review & Editing. **C.M.A.:** Methodology, Writing – Review & Editing. **C.H.S.:** Data curation, Software. **L.K.-W.:** Investigation, Data curation, Project administration. **B.A.M.:** Conceptualization, Software. **E.B.:** Investigation, Validation. **P.J.M.:** Writing - Review & Editing. **C.E.B.:** Conceptualization, Methodology, Writing – Review & Editing, Supervision, Funding acquisition, Resources.

