## Supplementary Material for "Convergent and Divergent Cerebellar Alterations in 22q11.2 Copy Number Variants"

**Supplementary Material for:**
Convergent and Divergent Cerebellar Alterations in 22q11.2 Copy Number Variants

Hoki Fung, Kathleen P. O'Hora, Rune Boen, Carolyn M. Amir, Charles H. Schleifer, Leila Kushan-Wells, Blake A. Miranda, Elizabeth Bondy, Paul J. Mathews, Carrie E. Bearden*

### SUPPLEMENTARY TABLES

#### Table S1. Full model output for total cerebellar volume permutation analysis

| Term | Estimate (B) | SE | Statistic | df | p-value | CI (2.5%) | CI (97.5%) |
|---|---|---|---|---|---|---|---|
| (Intercept) | 89467.99 | 5326.20 | 16.80 | 491.26 | <0.001 | 79003.05 | 99932.93 |
| Group_22qDel | -16072.03 | 1445.74 | -11.12 | 320.79 | <0.001 | -18916.36 | -13227.71 |
| Group_22qDup | 1579.76 | 2134.78 | 0.74 | 312.38 | 0.46 | -2620.60 | 5780.13 |
| Sex | 9423.80 | 1425.96 | 6.61 | 364.04 | <0.001 | 6619.64 | 12227.96 |
| Age (centered) | 53.98 | 77.73 | 0.69 | 482.39 | 0.49 | -98.75 | 206.71 |
| $Age^2$ | -11.22 | 4.23 | -2.65 | 507.00 | 0.01 | -19.53 | -2.91 |
| eTIV | 0.03 | 0.00 | 8.67 | 481.00 | <0.001 | 0.02 | 0.04 |

Linear mixed-effects model results for total cerebellar volume. The model includes fixed effects for group (TD, 22qDel, 22qDup), sex, age (mean-centered; linear and quadratic), and estimated intracranial volume (eTIV), with subject-level random intercepts.

**Table S2. Full model results for permutation-based regional cerebellar volume analyses**

| | 22qDel vs. TD | | | | | 22qDup vs. TD | | | | |
|---|---|---|---|---|---|---|---|---|---|---|
| **Region** | **B** | **SE** | **Statistic** | **p-value** | **FDR q** | **B** | **SE** | **Statistic** | **p-value** | **FDR q** |
| Corpus Medullare | -2268.34 | 264.22 | -8.59 | <0.001 | <0.001 | -169.75 | 387.93 | -0.44 | 0.66 | 0.73 |
| Left Lobules I-III | -59.80 | 27.37 | -2.18 | 0.03 | 0.03 | 10.67 | 40.17 | 0.27 | 0.79 | 0.79 |
| Right Lobules I-III | -99.59 | 26.41 | -3.77 | <0.001 | <0.001 | 22.34 | 38.75 | 0.58 | 0.56 | 0.73 |
| Left Lobule IV | -46.63 | 73.08 | -0.64 | 0.52 | 0.54 | 182.88 | 107.33 | 1.70 | 0.09 | 0.42 |
| Right Lobule IV | -107.65 | 75.05 | -1.43 | 0.15 | 0.16 | 49.99 | 110.28 | 0.45 | 0.65 | 0.73 |
| Left Lobule V | 1.82 | 63.51 | 0.03 | 0.98 | 0.98 | 90.78 | 93.21 | 0.97 | 0.33 | 0.61 |
| Right Lobule V | -275.43 | 62.15 | -4.43 | <0.001 | <0.001 | -71.09 | 91.30 | -0.78 | 0.44 | 0.68 |
| Vermis VI | -180.60 | 29.57 | -6.11 | <0.001 | <0.001 | -58.77 | 43.47 | -1.35 | 0.18 | 0.47 |
| Left Lobule VI | -922.26 | 165.20 | -5.58 | <0.001 | <0.001 | 99.97 | 242.98 | 0.41 | 0.68 | 0.73 |
| Right Lobule VI | -823.46 | 167.61 | -4.91 | <0.001 | <0.001 | 87.32 | 247.29 | 0.35 | 0.72 | 0.75 |
| Vermis VII | -124.98 | 20.54 | -6.08 | <0.001 | <0.001 | -100.05 | 30.21 | -3.31 | <0.001 | 0.03 |
| Left Crus I | -1559.62 | 239.09 | -6.52 | <0.001 | <0.001 | 243.62 | 351.65 | 0.69 | 0.49 | 0.72 |
| Left Crus II | -1082.77 | 173.38 | -6.25 | <0.001 | <0.001 | 138.72 | 254.67 | 0.54 | 0.59 | 0.73 |
| Left Lobule VIIB | -1178.50 | 152.14 | -7.75 | <0.001 | <0.001 | -102.85 | 223.61 | -0.46 | 0.65 | 0.73 |
| Right Crus I | -1194.20 | 227.38 | -5.25 | <0.001 | <0.001 | 508.82 | 335.08 | 1.52 | 0.13 | 0.47 |
| Right Crus II | -927.54 | 159.77 | -5.81 | <0.001 | <0.001 | 300.52 | 234.84 | 1.28 | 0.20 | 0.47 |
| Right Lobule VIIB | -850.42 | 157.77 | -5.39 | <0.001 | <0.001 | 298.59 | 231.72 | 1.29 | 0.20 | 0.47 |
| Vermis VIII | -132.02 | 39.02 | -3.38 | <0.001 | <0.001 | 112.50 | 57.54 | 1.96 | 0.05 | 0.36 |
| Left Lobule VIIIA | -923.42 | 154.92 | -5.96 | <0.001 | <0.001 | -213.28 | 227.44 | -0.94 | 0.35 | 0.61 |
| Left Lobule VIIIB | -527.12 | 91.40 | -5.77 | <0.001 | <0.001 | 133.60 | 134.19 | 1.00 | 0.32 | 0.61 |
| Right Lobule VIIIA | -812.62 | 122.64 | -6.63 | <0.001 | <0.001 | -358.09 | 179.97 | -1.99 | 0.05 | 0.36 |
| Right Lobule VIIIB | -544.22 | 85.26 | -6.38 | <0.001 | <0.001 | -178.91 | 125.12 | -1.43 | 0.15 | 0.47 |

| | | | | | | | | | |
|---|---|---|---|---|---|---|---|---|---|
| Vermis IX | -168.46 | 20.20 | -8.34 | <0.001 | <0.001 | 61.02 | 29.69 | 2.06 | 0.04 | 0.36 |
| Left Lobule IX | -389.11 | 87.22 | -4.46 | <0.001 | <0.001 | 131.92 | 128.33 | 1.03 | 0.30 | 0.61 |
| Right Lobule IX | -319.26 | 86.88 | -3.67 | <0.001 | <0.001 | 236.69 | 127.74 | 1.85 | 0.06 | 0.36 |
| Vermis X | -22.88 | 7.41 | -3.09 | <0.001 | <0.001 | -6.02 | 10.90 | -0.55 | 0.58 | 0.73 |
| Left Lobule X | -24.71 | 9.89 | -2.50 | 0.01 | 0.02 | -18.50 | 14.52 | -1.27 | 0.20 | 0.47 |
| Right Lobule X | -28.50 | 11.28 | -2.53 | 0.01 | 0.01 | -13.93 | 16.56 | -0.84 | 0.40 | 0.66 |

Results from linear mixed-effects models assessing group differences across 28 anatomically defined cerebellar subregions. Each model included fixed effects for sex, age (mean-centered; linear and quadratic), estimated intracranial volume (eTIV), and group (TD, 22qDel, 22qDup), with a subject-level random intercept to account for repeated measures. In these models, TD controls served as the reference group, and results are reported for 22qDel vs. TD (left) and 22qDup vs. TD (right) comparisons.

**Table S3. Full model results for permutation-based regional cerebellar volume analyses (22qDel as reference; 22qDup vs. 22qDel)**

| | 22qDup vs. 22qDel | | | | | | | |
|---|---|---|---|---|---|---|---|---|
| **Region** | **Estimate (B)** | **SE** | **Statistic** | **df** | **CI (2.5%)** | **CI (97.5%)** | **p-value** | **FDR q** |
| Corpus Medullare | 2098.59 | 403.06 | 5.21 | 314.89 | 1305.56 | 2891.61 | <0.001 | <0.001 |
| Left Lobules I-III | 70.47 | 41.63 | 1.69 | 278.23 | -11.47 | 152.41 | 0.09 | 0.12 |
| Right Lobules I-III | 121.93 | 40.20 | 3.03 | 284.78 | 42.80 | 201.06 | <0.001 | <0.001 |
| Left Lobule IV | 229.50 | 111.56 | 2.06 | 304.24 | 9.97 | 449.04 | 0.04 | 0.05 |
| Right Lobule IV | 157.64 | 114.68 | 1.37 | 301.01 | -68.04 | 383.32 | 0.17 | 0.20 |
| Left Lobule V | 88.96 | 96.74 | 0.92 | 289.11 | -101.44 | 279.35 | 0.36 | 0.40 |
| Right Lobule V | 204.34 | 94.93 | 2.15 | 298.64 | 17.54 | 391.15 | 0.03 | 0.05 |
| Vermis VI | 121.83 | 45.21 | 2.69 | 295.91 | 32.85 | 210.81 | 0.01 | 0.01 |
| Left Lobule VI | 1022.22 | 252.83 | 4.04 | 301.62 | 524.70 | 1519.75 | <0.001 | <0.001 |
| Right Lobule VI | 910.78 | 257.53 | 3.54 | 297.33 | 403.97 | 1417.58 | <0.001 | <0.001 |
| Vermis VII | 24.93 | 31.43 | 0.79 | 303.24 | -36.91 | 86.78 | 0.43 | 0.44 |
| Left Crus I | 1803.24 | 365.88 | 4.93 | 308.30 | 1083.31 | 2523.17 | <0.001 | <0.001 |
| Left Crus II | 1221.49 | 264.75 | 4.61 | 300.50 | 700.48 | 1742.50 | <0.001 | <0.001 |
| Left Lobule VIIB | 1075.66 | 232.57 | 4.63 | 293.17 | 617.93 | 1533.38 | <0.001 | <0.001 |
| Right Crus I | 1703.01 | 348.85 | 4.88 | 311.67 | 1016.61 | 2389.42 | <0.001 | <0.001 |
| Right Crus II | 1228.06 | 244.26 | 5.03 | 303.46 | 747.41 | 1708.72 | <0.001 | <0.001 |
| Right Lobule VIIB | 1149.01 | 240.85 | 4.77 | 304.98 | 675.06 | 1622.95 | <0.001 | <0.001 |
| Vermis VIII | 244.53 | 59.91 | 4.08 | 315.19 | 126.65 | 362.40 | <0.001 | <0.001 |
| Left Lobule VIIIA | 710.14 | 236.26 | 3.01 | 307.68 | 245.24 | 1175.04 | <0.001 | 0.01 |
| Left Lobule VIIIB | 660.72 | 139.40 | 4.74 | 310.76 | 386.42 | 935.02 | <0.001 | <0.001 |
| Right Lobule VIIIA | 454.53 | 186.64 | 2.44 | 308.03 | 87.28 | 821.78 | 0.02 | 0.02 |
| Right Lobule VIIIB | 365.31 | 129.73 | 2.82 | 317.27 | 110.08 | 620.54 | 0.01 | 0.01 |
| Vermis IX | 229.48 | 30.87 | 7.43 | 309.70 | 168.73 | 290.23 | <0.001 | <0.001 |
| Left Lobule IX | 521.03 | 133.54 | 3.90 | 310.07 | 258.27 | 783.78 | <0.001 | <0.001 |
| Right Lobule IX | 555.95 | 132.89 | 4.18 | 308.32 | 294.46 | 817.44 | <0.001 | <0.001 |
| Vermis X | 16.86 | 11.34 | 1.49 | 315.74 | -5.44 | 39.16 | 0.14 | 0.17 |
| Left Lobule X | 6.20 | 15.05 | 0.41 | 303.64 | -23.42 | 35.83 | 0.68 | 0.68 |

| | | | | | | | | |
|---|---|---|---|---|---|---|---|---|
| Right Lobule X | 14.57 | 17.18 | 0.85 | 321.72 | -19.23 | 48.38 | 0.40 | 0.43 |

Results from linear mixed-effects models assessing group differences across 28 anatomically defined cerebellar subregions. Each model included fixed effects for sex, age (mean-centered; linear and quadratic), estimated intracranial volume (eTIV), and group (TD, 22qDel, 22qDup), with a subject-level random intercept to account for repeated measures. In these models, 22qDel was the reference group, and results for the 22qDup vs. 22qDel comparison are reported here.

### SUPPLEMENTARY FIGURES

**Figure S1. Group differences in total and regional cerebellar volumes.**

Standardized beta coefficients and 95% confidence intervals are shown for the 22qDup group relative to the 22qDel group. Results were derived from linear mixed-effects models controlling for sex, mean-centered age (linear and quadratic), and estimated intracranial volume (eTIV), with a subject-level random intercept to account for repeated measures. Asterisks indicate significance after FDR correction for regional volumes: q < .05 (*), q < .01 (**), q < .001 (***). TOTAL refers to total cerebellar volume and is shown with uncorrected p-value, as it was not included in the FDR correction.

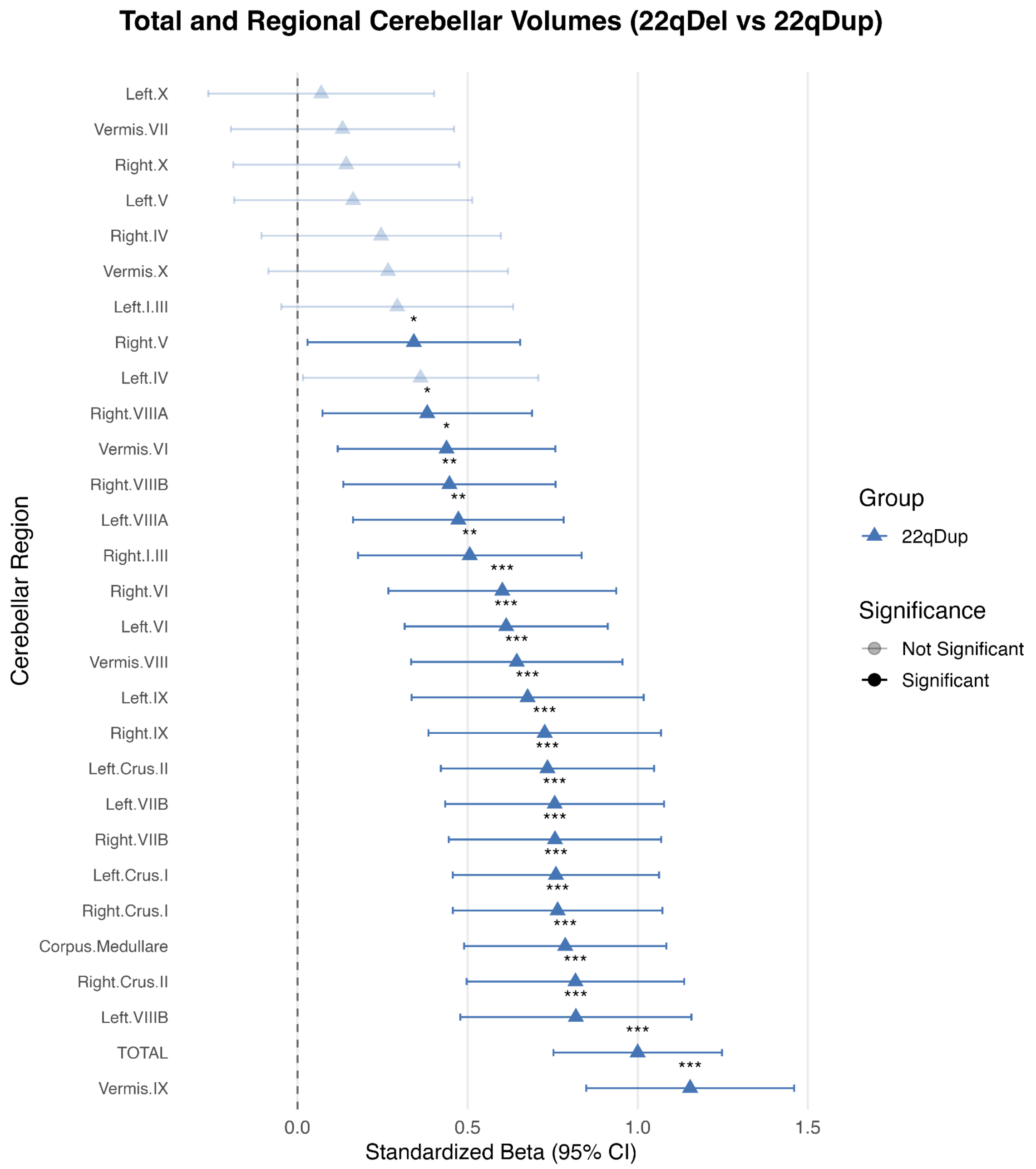
